# K-mer Motif Multinomial Mixtures, a scalable framework for multiple motif discovery

**DOI:** 10.1101/096735

**Authors:** Brian L. Trippe, Sandhya Prabhakaran, Harmen J. Bussemaker

**Affiliations:** Department of Biological Sciences, Columbia University, New York, 10027, USA; Department of Systems Biology, Columbia University Medical Center, New York, 10032, USA

## Abstract

1

**Motivation:** The advent of inexpensive high-throughput sequencing (HTS) places new demands on motif discovery algorithms. To confront the challenges and embrace the opportunities presented by the growing wealth of information tied up in HTS datasets, we developed K-mer motif multinomial mixtures (KMMMs), a flexible class of Bayesian models for identifying multiple motifs in sequence sets using K-mer tables. Advantages of this framework are inference with time and space complexities that only scale with K, and the ability to be incorporated into larger Bayesian models.

**Results:** We derived a class of probabilistic models of K-mer tables generated from sequence containing multiple motifs. KMMMs model the K-mer table as a multinomial mixture, with motif and background components, which are distributions over K-mers overlapping with each of the latent motifs and over K-mers that do not overlap with any motif, respectively. The framework casts motif discovery as a posterior inference problem, and we present several approximate inference methods that provide accurate reconstructions of motifs in synthetic data. Finally we apply the method to discover motifs in DNAse hypersensitive sites and ChIP-seq peaks obtained from the ENCODE project.

## 2 Introduction

The efficient identification of multiple motifs within large DNA sequences is an important problem in modern genomics. With the realization of inexpensive high-throughput DNA sequencing (HTS) we are confronted with large sets of sequences that we are interested in characterizing and comparing against one another. For example, in the realm of regulatory genomics, several experimental methods allow us to probe many aspects of regulatory state across entire genomes. In comparative analyses, where we may want to consider these aspects over many samples, the datasets quickly amount to many billions of bases of sequenced DNA [Encode, 2012, Kundaje *et al*, 2015]. In this context and in others, we may want to assess the motifs present in large sequence sets, as these patterns are a primary functionally relevant characteristic of genomic sequence.

A wide range of approaches to discovering motifs have been developed over the past two decades, dating back to the MEME [Bailey and Elkan, 1994]. Since MEME, there have been many other methods which take different approaches. For a more in depth review, see Das and Dai 2007[Das and Dai, 2007]. As next generation sequencing has lead to increasingly large sequence sets, a variety of more recent methods have been developed, many focusing on ChIP-Seq including MEME-ChIP, an adaptation of MEME tailored towards ChIP-Seq, HOMER and others[Heinz *et al*, 2010, Machanick and Bailey, 2011, Tran *et al*, 2014]. Many effective recent methods have been discriminative, working off of an objective of distinguishing peaks from background sequence, some of which have used K-mer features [Arvey *et al*, 2012, Setty *et al*, 2015].

KMMMs stand in contrast to these methods as a generative model of K-mer tables constructed from sequences containing motifs. Motifs and mixing proportions are modeled as latent variables and are learned by posterior inference. Whereas discriminative methods identifying motifs from ChIP-Seq aim to classify sequences as either bound or unbound, KMMMs aim to learn a statistical summary of a single sequence set based on the motifs present within it. KMMMs combine the strength of generative models such as MEME with representational efficiency of K-mer based approaches.

In addition to concerns for efficiency of motif discovery, when trying to better understand complex biological processes we often want to integrate multiple forms of data together with our prior knowledge about the relationships between them. This is particularly true in the context of regulatory genomics. Bayesian statistics and graphical models present a useful framework for such tasks and allow one to investigate the posterior distributions over large sets of latent variables that represent biologically meaningful quantities. Common methods of Bayesian inference require iterating over these latent variables and performing computation with respect to the others. For example, in coordinate ascent variational inference we iterate over latent variables, updating variational approximations of their posteriors until convergence and in Gibbs Sampling we iteratively sample from complete conditional distributions to obtain samples from these posteriors. For this reason, when we are working with large datasets of sequences in the context of a larger Bayesian model, the use of algorithms whose run-times scale with the quantity of the sequence quickly becomes intractable.

We introduce K-mer motif multinomial mixtures (KMMMs), a Bayesian framework for discovering multiple motifs that overcomes these limitations of earlier models. The value of the KMMM is twofold. First it provides an effective and flexible method for analyzing sequence composition and performing *de novo* discovery of multiple motifs simultaneously; it is generalized over a range of motif and background models, with specific instances of a KMMM being defined by these choices. Second, as a multinomial mixture model over the K-mer table, space and time complexities of inference do not scale with the quantity of sequence; only the preprocessing step of creating this table takes linear time. While the complexity does scale exponentially with the size of K-mers, a relatively modest value of *K* is sufficient, as the size of the inferred motifs is not limited by it. KMMMs thereby provide a scalable framework that can be easily expanded upon and integrated into larger Bayesian models.

## 3 Approach: motifs and K-mer tables

KMMMs define a generative model in which motifs are modeled as latent variables and define a distribution over the K-mer table which is modeled as an observed variable. To discover motifs, sequences are first collapsed into a K-mer table (i.e., a table of counts of *K* base long subsequences that occur within a larger sequence set), and motifs are found by posterior inference (Figure 1). Before formally introducing KMMMs, we motivate them with and discussion of motifs and K-mer tables in sections 3.1 and 3.2.

**Figure 1:**
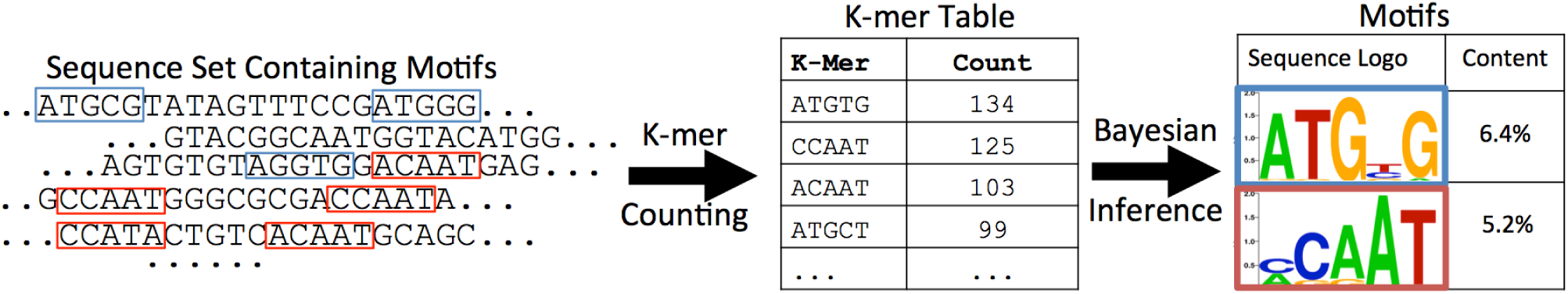
Schematic of motif discovery with the KMMM. First, a sequence set is collapsed into a K-mer table using a sliding window. Motifs are then found by posterior inference.

### 3.1 Motifs

A DNA sequence motif defines a distribution over sequences of some length (typically 5 to 20 bases) that occur repeatedly in a larger DNA sequence or set of sequences [Das and Dai, 2007]. For example, a number of regulatory motifs occur repeatedly across the promoters of multiple genes and enable modulation of expression through their interaction with transcription factors that have affinity for these sequences.

A common method of representing motifs which we use in this model approximates a length *L* motif as a sequence of *L* independent distributions over the 4 bases (A, C, T and G) at each position. As such, we define a length *L* motif, *β* = (*β*_1_, *β*_2_,…, *β_L_*), as a sequence of categorical distributions over bases, where each *β_l_* defines the probabilities of seeing each of the nucleotides at position *l*. For a sequence, *s* = (*s*_1_, *s*_2_,…, *s_L_*), associated with the motif, each base, *s_l_*, is drawn from the distribution *β_l_*. Accordingly, the probability of the sequence can thus be decomposed into a product of the mononucleotide probabilities:

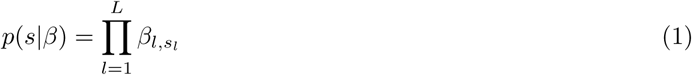

This common approximate representation of sequence preference is often referred to as a position-specific frequency matrix or position weight matrix (PWM).

### 3.2 K-mer tables

A K-mer table may be generated by sliding a length *K* window along a sequence, incrementing the count of the windowed sequence at each position. This abandons all information about the order of the sub-sequences as well as their context in the larger sequence. This way of working with a K-mer table representation of sequences is akin to the ‘bag-of-words’ construction used widely in document modeling [Aldous, 1985, Blei *et al*, 2003]. Considering K-mer tables to confront the challenges of modeling motifs in large sequence sets is reasonable in view of two observations about the nature of K-mer tables and the motif discovery problem. First, the information content of motifs is purely in local context (i.e. the co-occurrence probabilities of bases within the motif) while the global position carries no information. Second, the space and time complexities of operations on a K-mer table scale with *K* (the length of the K-mers) rather than with the sequence length. This is a valuable property when we are handling large quantities of sequence data. On the basis of these two observations, by performing inference with K-mer tables, we should be able to identify motifs with minimal loss of precision and with time complexity that does not scale with sequence length. Recently, K-mer tables have been used for efficient quantification of RNA-seq data, another problem in which the scaling properties of typical alignment based algorithms make their applications to large sequence sets unwieldy [Bray *et al*, 2016].

Intuition into our approach can be gained by considering how the presence of a motif in sequence will impact distribution of counts of subsequences. To put this in another way, how does a motif component distribute its mass across the 4^*K*^ dimensional discrete space of the K-mer table? As an example, consider how the distribution of 4-mers is impacted by the presence of a motif that has probability 1.0 of being “ACCG” and probability of 0. 0 of being any other 4-mer. Were we to generate sequence by repeatedly appending either an ‘A’, ‘C’, ‘T’, or ‘G’ with equal probabilities, we would expect the distribution over the table to be roughly uniform. However, if with some probability we instead sample from our motif (and append “ACCG”), this distribution will shift. The details of the generative model by which this sequence was produced are explained precisely in the supplement. A visualization of this K-mer table as a heatmap is provided in Figure 2A. As expected, increased density is observed for the subsequence “ACCG”. However, we additionally see higher density on 4-mers ending in “ACC” (i.e “AACC”, “CACC”, “GACC” and “TACC”) and slightly increased density on sequences ending with “AC” (Figure 2A, column 12), *i.e*., sequences that overlap partially with the motif (Figure 2B). In this way a motif exerts a unique ‘fingerprint’ on the K-mer table.

**Figure 2:**
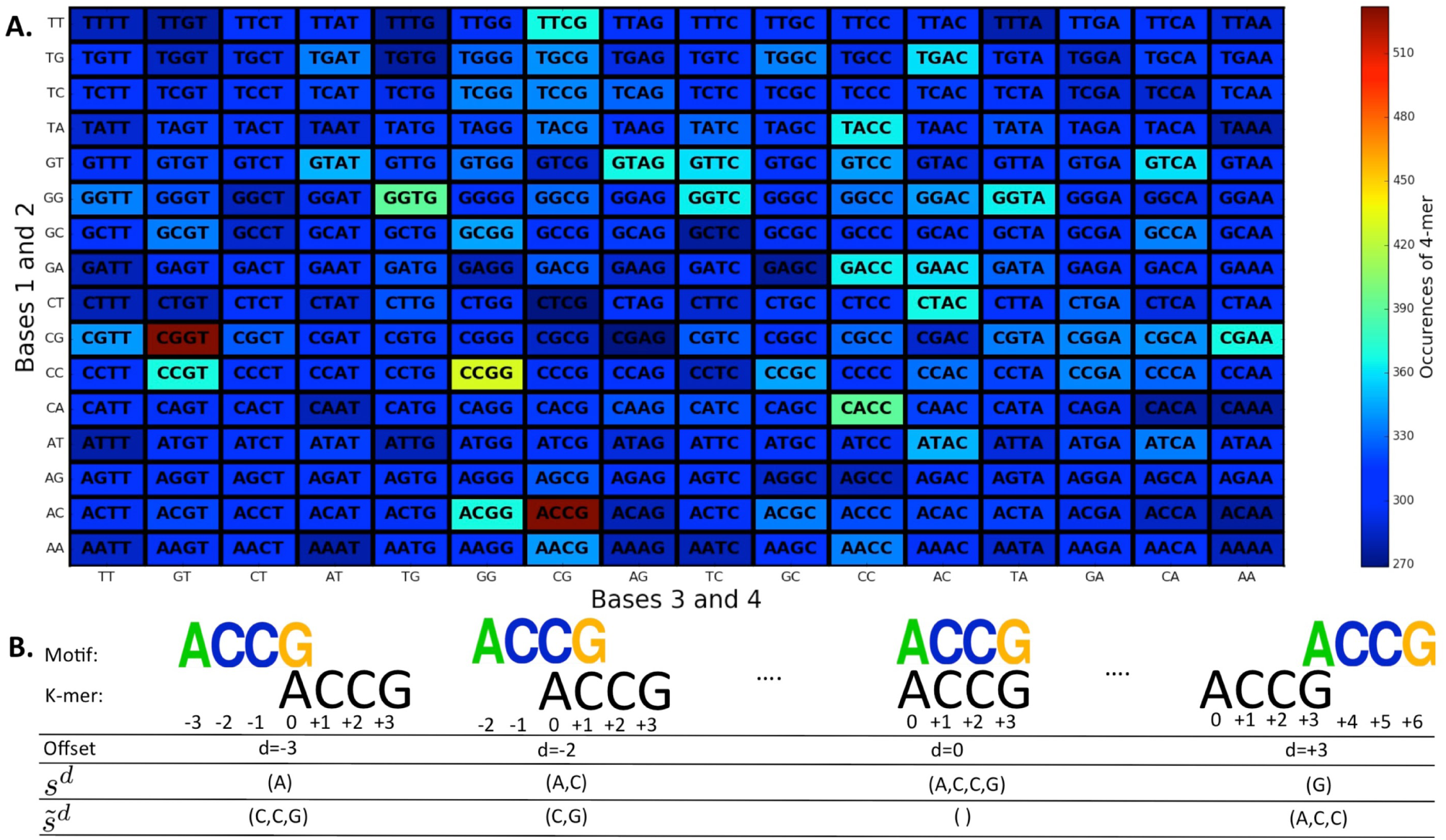
Sequences of arbitrary length can be represented with K-mer tables, counts of all length *K* subsequences. When sequences contain patterns, certain K-mers will occur more often than others. A.) Heatmap of counts of a 4-mers (*K* = 4) in one megabase of synthetic sequence generated by adding single bases with equal probability with additional occurrences of the sequence ‘ACCG’. B.) A K-mer can overlap with an occurrence of a sequence motif with one of several possible offsets. Therefore, a K-mer that aligns with a motif with a partial overlap, *s^d^*, with part of the K-mer arising from background, 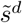 may still have an increased count.

With this observation in mind, we move to consider the counts in a K-mer table generated from sequence with multiple motifs as a draw from a multinomial distribution. The probability of each sequence is the sum of its probabilities of overlapping with each of the motifs present in the underlying sequence and of occurring in the background distribution. By making the assumption that motifs are distributed sparsely enough that they never overlap with the same K-mer window, we are able to decompose this larger multinomial distribution into a mixture of multinomial components. In this mixture the components are: (i) motifs components, which each comprise the set of sequences that overlap with an instance of that motif (either entirely or with a partial overlap) and (ii) a background component, comprised of sequences not overlapping with a motif. In this framing, a K-mer table constructed from sequence with *M* motifs will have *M* + 1 mixture components. This perspective on the composition of sequences naturally leads to the construction of K-mer Motif Multinomial Mixtures (KMMMs).

## 4 K-mer motif multinomial mixtures

We formally introduce K-mer motif multinomial mixtures in this section by deriving the likelihood distribution over the K-mer table, which we denote as *X*, induced by a sequence segmentation model in which stochastic words are used [Bailey and Elkan, 1994, Bussemaker *et al*, 2000, Gupta and Liu, 2011]^1^. We first show that the likelihood takes the form of a multinomial mixture, thereby formally introducing KMMMs. Next, we cast the KMMM in a Bayesian light and provide a complete generative model and graphical model representation.

For clarity and simplicity, we first introduce KMMMs with several simplifications that reduce the complexity of notation needed. These include: ignoring reverse complementary of sequences, restricting the length of motifs to the length of the K-mers, assuming we are provided with an accurate model of background sequence, assuming that the number of motifs and their lengths are known a priori. These assumptions are subsequently lifted in 5.3 and in the supplement.

### 4.1 Decomposing the likelihood of the K-mer table into a mixture of multinomials

We consider the likelihood of the counts in the K-mer table, *X*, given a set of *M* motifs, *β* = (*β*_1_, *β*_2_,…,*β_M_*), a model of background sequence, *ϕ*, and mixing proportions, *γ* = (*γ*_0_, *γ*_1_,…, *γ_M_*) where *γ*_0_ refers to the portion of the background component. Because a K-mer table is counts in a discrete space, we can model it as a draw from a multinomial distribution:

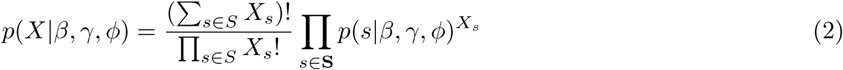

where *S* = {*A*, *C*, *T*, *G*}^*K*^ and for each *s* ∈ *S*, *X_s_* is the number of occurrences of *s*.

From here, we expand the probability of observing a K-mer given our latent variables, *p*(*s*|*β*, *γ*, *ϕ*). We make the assumption that each K-mer must either come entirely from background or overlap with exactly one of the *M* motifs. While this assumption is broken when motifs are very closely packed (allowing a single K-mer to overlap with more than one motif), this approximation will be close to reality when motifs are distributed sparsely. This key assumption allows us to expand out this likelihood by marginalizing over possibilities:

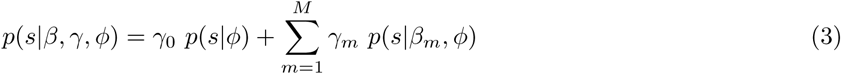

where *β_m_* is the PWM representation of motif *m* and *γ*_0_ is the mixing proportion of the background component.

It is in this step that our model becomes a multinomial mixture model. Each term in the expression above defines a mixture component which is itself a multinomial distribution over the K-mer table. Here we see an intuitive interpretation of the mixing proportions, *γ*. *γ*_0_ represents the portion of K-mers arising entirely from background, and for each *m* ≠ 0, *γ_m_* represents the portion of K-mers overlapping with an occurrence of motif *m*.

All that now remains is to formally define the probabilities of sequences in each of these components. The background model, *ϕ*, directly provides the first term, *p*(*s*|*ϕ*), but we need to do a bit more work for the motif components, *p*(*s*|*β_m_*, *ϕ*). Here, we again use marginalization, in this case over the possible offsets, *d*^2^:

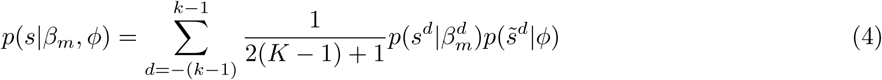

For each of the possible offsets, we posit that the probability of the K-mer is the product of the probabilities of observing the segment overlapping with the motif, *s^d^*, under the overlapping portion of the motif, 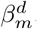 and the probability of observing the rest of the sequence, 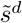, from the background model (Figure 2B). This fully expands the multinomial mixture, breaking each of the motif components into 2(*K* – 1) + 1 sub-components of equal weight^3^, each defining a multinomial distribution over the K-mer table.

This completes the definition of KMMMs. The distribution over K-mer counts in a sequence containing *M* motifs is modeled as a mixture of 1 + *M*(2(*K* – 1) + 1) multinomial components: one background component with a mixing proportion *γ*_0_, and, for each motif *m*, 2(*K* – 1) + 1 components corresponding to different offsets, each with a mixing proportion *γ_m_*/[2(*K* – 1) + 1].

In the case of the PWM motif model, when motifs are the same length as the K-mers, the likelihood of a K-mer given each of these offsets can be explicitly formulated as

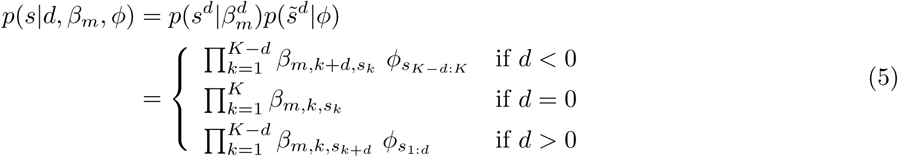

We emphasize, however, that KMMMs are not restricted to this underlying motif model. Motif models that capture higher order characteristics of motifs (*e.g.* dinucleotide preferences) can be used as well. We detail these in section 11.1.

### 4.2 A generative probabilistic model of K-mer tables

We now take a Bayesian perspective. We explicitly model the motifs, mixing proportions and component and offsets assignments as latent variables and we propose prior distributions over them. Since the mixing proportions, *γ*, lie on the simplex, we place on them a Dirichlet prior *α*. Each of the per-position distributions over bases that define the motifs, *β_m,k_*, also lie on the simplex, allowing us to place a uniform Dirichlet prior, π, upon them as well. Lastly, we place a uniform multinomial prior over the offset assignments for K-mers assigned to motif components, *τ*.

Provided with these priors and the formulation of the likelihood in equations 2 through 4, we present the following generative process for producing a list of observed K-mers, *s*, and the corresponding graphical model in Figure 3:

1. Choose each of *K* positions of the *M* motifs, *β_m,k_* ~ Dir(*π*)
2. Choose mixing proportions, *γ* ~Dir(*α*).
3. For each of the *N* K-mers *s_n_*:
  a. Choose a component *m_n_* ~ Mult(*γ*)
  b. If *m_n_* = 0, draw the K-mer from the background component
    i. Choose *s_n_* ~ Mult(s_n_|*ϕ*)
  c. If not *m_n_* = 0, draw the K-mer from motif component, *m_n_*
    i. Choose an offset *d_n_* ~ Uniform(–(k – 1), –(k – 2),…, (k — 1)|*τ*)
    ii. Choose 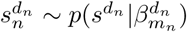
    iii. Choose 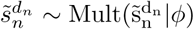
    iv. *s_n_* 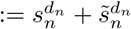

**Figure 3:**
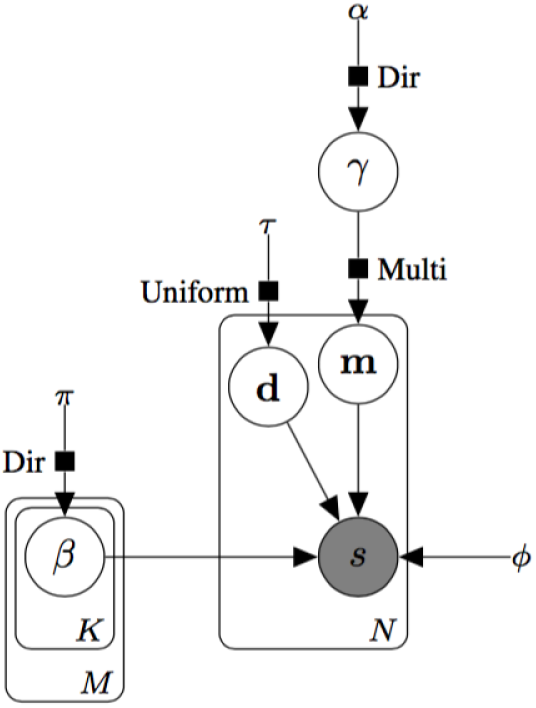
Graphical model representation of a KMMM. The plate on the right side represents the multiplicity of *N* K-mer subsequences.

With this construction, we can calculate the probability of the full joint distribution over the latent variables and observed K-mers using the chain rule and knowledge of the conditional independences of the model. We separate the latent variables and the observed K-mers, and factorize them into the individual distributions that compose our motif representation:

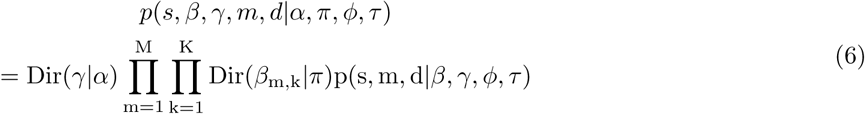

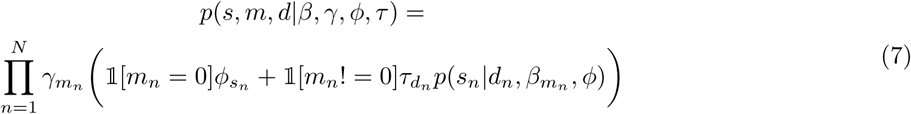

This model specification is akin to the ‘Bag of Words’ model in document modeling; it defines the probability of one permutation of the set of K-mers and does not yet directly correspond to the distribution over the K-mer table. To make this connection, we collapse the observed list of K-mers, *s*, into a K-mer table, *X*, and collapse the associated latent component assignments, *m* and *d*, into a new latent variable, *z*, where *z_s,m,d_* is the number of occurrences of the K-mer, *s*, assigned to motif *m* with offset *d*. We thus move from considering the distribution over permutations of a set of K-mers and component assignments to the distribution over combinations of these variables and in this way we transition from a ‘Bag of Words’ model to a multinomial model. With this new notation, we can rewrite equation 7 as:

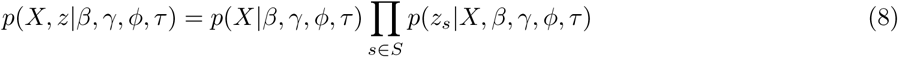

Where the first term of the right hand side is the multinomial probability of the K-mer table derived in section 4.1 and the factors in the second term are multinomials over the component assignments of the occurrences of each of the K-mers. These factors expand out as:

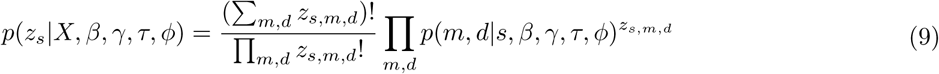

where ∀*s* ∈ *S*, Σ_*md*_ *z_s,m,d_* = *X_s_*, and the probabilities of each of the component assignments are found by normalizing the likelihoods given by:

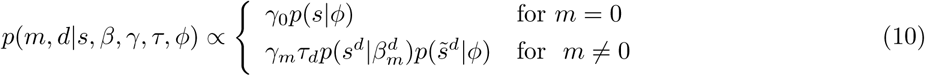

By considering the counts of each K-mer attributed to each of the different mixture components, rather than each of the individual subsequences, our sequence representation no longer scales in size with sequence length.

## 5 Posterior inference

This section discusses posterior inference KMMMs. To be precise, we are interested in inferring the motifs, *β*, their mixing proportions, *γ*, and component assignments, *z*, given their priors and the K-mer table, *X*:

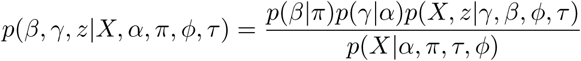

Direct posterior computation is intractable due to the evidence term, *p*(*X*|*α*,π, *ϕ*,*τ*), as is frequently the case for latent variable models. In this model, calculation of the evidence would require integrating over all possible motifs, their proportions and possible partitions of the K-mer table across the mixture components. To confront this challenge we turn to approximate inference methods.

Approximate inference is straight-forward in KMMMs because it is a conditionally conjugate model. In this section, we first demonstrate this property in the simplest instance of a KMMM, using a PWM motif model and a fixed random background model. We then briefly discuss inference algorithms and extensions of the model which enable motif discovery in a realistic setting while maintaining the model’s conditional conjugacy.

### 5.1 Conditional conjugacy in KMMMs

The property of conditional conjugacy in latent variable models provides that the complete conditional distribution over each of the model’s latent variables (i.e. its conditional distribution given its Markov blanket) falls in the same family as its prior distribution. In conditionally conjugate models, both sampling and variational methods are easier to apply because sampling distributions and coordinate updates, respectively, have an analytic, closed form [Beal, 2003, Bishop 2006]. In this section, we show that the distributions over the three latent variables of KMMMs, *β*, *γ* and *z* are conditionally conjugate.

For the mixing proportions of the motif and background components, *γ*, the complete conditional takes the following form^4^:

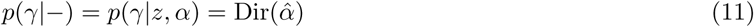

where:

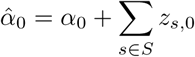

and for *m* ≠ 0:

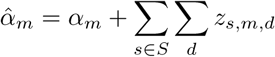

This closed form relies on Dirichlet-multinomial conjugacy, which guarantees that posterior over a latent multinomial random variable drawn from a known Dirichlet prior and conditioned on samples drawn from that multinomial, takes the form of a Dirichlet.

Our ability to formulate the complete conditional over motif parameters relies on Dirichlet-multinomial conjugacy as well. In this case, given the Dirichlet prior, π, and the observed bases at each position, the complete conditionals over each *β*_*m*,*k*_ are also Dirichlet distributed. However, directly formulating these distributions requires additional accounting, as the base counts are not encoded in the structure of the latent variable as directly as the component assignments.

To reason about this more precisely, we introduce an additional piece of notation for each of the motifs, *η_m_*, which refers to the counts of bases from all of the alignments of K-mers against motif *m*. With this notation, *η_m,i,A_* is the number of times we have observed an ‘A’ at position *i* in motif *m*. The value of each component of *η* is defined deterministically as a function of the latent variable *z* through a mapping provided in Algorithm 2 in section 10.1.

With this new notation established, the form of our complete conditional is trivially:

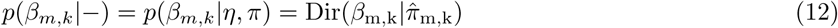

where:

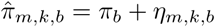

The remaining latent variable, *z*, also exhibits conditional conjugacy:

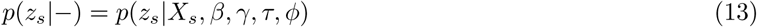

This fact is trivially true because the complete conditional is precisely the sampling distribution defined in equations 9 and 10.

### 5.2 Inference methods

Bayesian methods separate the problems of modeling and inference, i.e. the process of defining a generative latent variable model, as we did in section 4.2, is done separately from the design and implementation of a posterior inference algorithm. For KMMMs, the property of conditional conjugacy demonstrated in the previous section enables straightforward implementations of a variety of common inference algorithms. One common method is Gibbs sampling, in which we iteratively sample updates of each latent variable from the complete conditionals given in equations 11, 12 and 13 [Geman and Geman, 1984]. A second method is coordinate ascent mean-field variational inference, which can be implemented with closed form updates in conditionally conjugate model[Jordan *et al*, 1999, Blei *et al*, 2016]. Additionally, this property makes the model amenable to inference with stochastic variational inference [Hoffman *et al*, 2013]. We provide a more in depth explanation of these inference methods in sections 10.2, 10.3 and 10.4. A third option for inference is expectation maximization (EM). This procedure is closely related to variational inference, expectations of component assignments are updated in the E-step according to the complete conditional given in equation 13, and in the M step, the latent motif and mixing proportions are updated to maximize probability of the joint distribution given the expectations of all component assignments.

Conditional conjugacy additionally allows KMMMs to be used to quantify the presence of a known motif (or a set of motifs) when motifs of interest are known *a priori*. This is done by using alterations of the same procedures described above, in which the update step for motifs is skipped. KMMMs thereby allow quantification of motifs in arbitrarily large sequences (or collections of sequence) once a K-mer table is generated.

### 5.3 Several useful extensions of KMMMs maintain conditional conjugacy

The core model of KMMMs contains many restrictions which hinder its utility in real motif discovery problems. The actual genomic sequence between occurrences of motifs is more complex than the random background sequence used in our illustrative example in Figure 2; GC content and dinucleotide and trinucleotide frequencies can vary significantly from purely random sequence.

If these aspects of sequence are not taken into account, motifs that reflect the low complexity characteristics of background sequence may be learned (*e.g.* as a Poly-A motif). To confront this issue, we can model the distribution over background sequence as a Markov chain [Bussemaker *et al*, 2000]. Furthermore, to better distinguish complexities of background sequence from that of motifs, we consider the background model as an additional latent variable and fit it simultaneously with motifs. We discuss this extension in greater depth in section 11.2.

Additionally, the core model requires that the number of motifs and their lengths be known *a priori*, which is generally not the case. We eliminate these two requirements by introducing two Bayesian nonparametric extensions to KMMMs. First to answer the question of modeling sequence with an unknown number of motifs, we replace the finite Dirichlet prior over mixing proportions with a Dirichlet process prior, which allows inference over an infinite number of motif components to be performed by Gibbs sampling. To infer unknown motif lengths, we incorporate a latent shape variable and place a truncated geometric prior over motif lengths. Full details of these two extensions are included in sections 11.4 and 11.3.

## 6 Results

### 6.1 Known motifs are recovered from synthetic sequence data

To assess the ability of KMMMs and our inference methods to correctly identify motifs and their mixing proportions, we applied them to synthetic sequences generated according to Algorithm 1 in section 9.1 using motifs PWMs corresponding to mammalian transcription factors obtained from the JASPAR motif database using fixed random background model (i.e. with probability 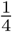 of each base not part of a motif) [Mathelier *et al*, 2013]. We present the results of two such experiments.

To test the ability of KMMMs to recover long motifs, we generated synthetic sequence using the 19-base long PWM for CTCF (JASPAR-ID: MA0139.1) and faithfully recovered it from an 8-mer table, with *R*^2^ = 0.9939 (Figure 4A and B). Next, to test its ability to recover multiple motifs from a mixture we generated sequence with two 8-base long motifs, Nobox(JASPAR-ID: MA0125.1) and Sox-17(JASPAR-ID: MA0078.1) and recovered them both from a single 7-mer table, *R*^2^ = 0.99995 (Figure 4C and D). In this test, we used the Bayesian nonparametric extension detailed in section 11.4 to correctly identify the number of latent motifs as part of inference.

**Figure 4:**
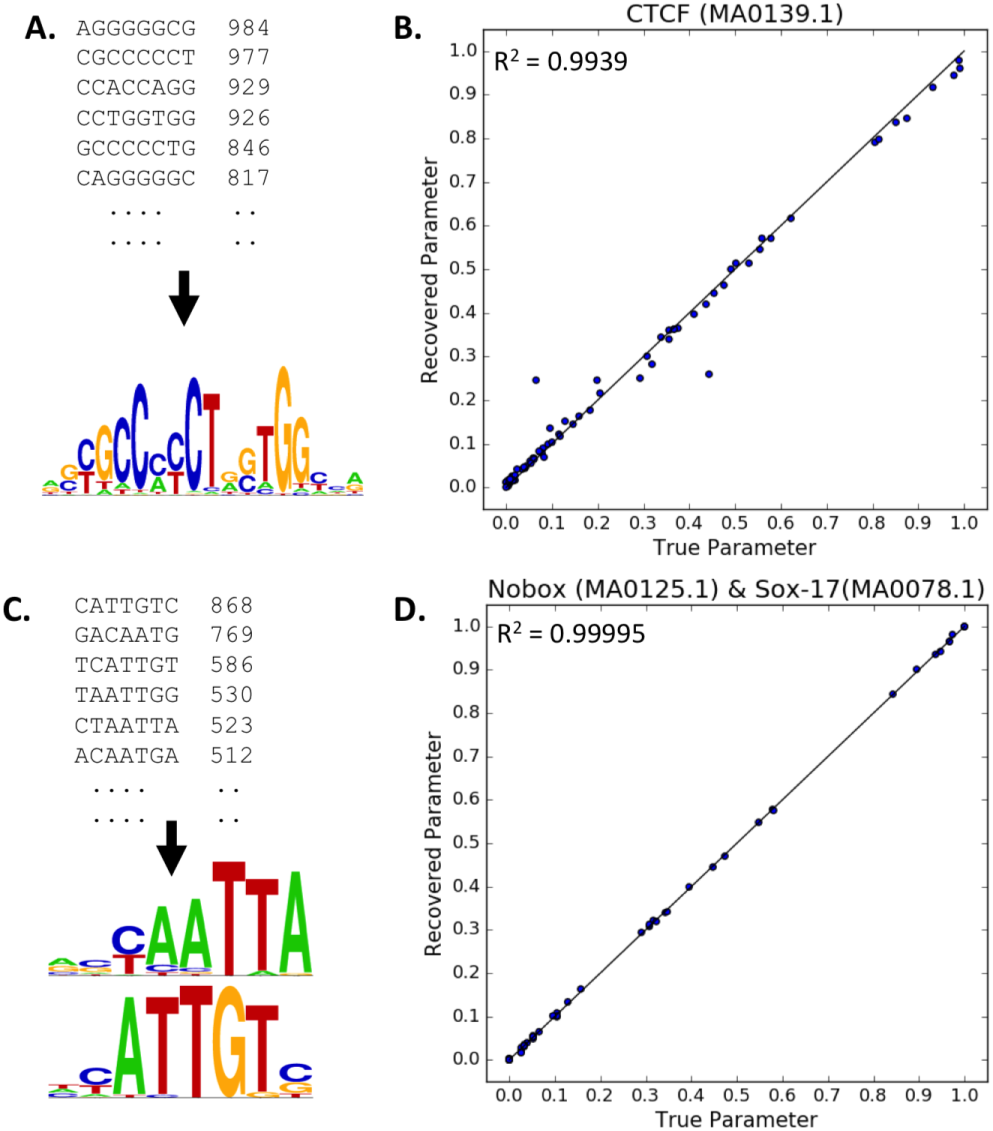
Motifs are faithfully recovered from K-mer tables in two applications of a KMMM to synthetic data using Gibbs sampling. **A.** The counts of top occurring 8-mers in a table constructed from synthetic sequence generated with a 19 base long CTCF motif (JASPAR ID MA0139.1) and a logo representing the motif recovered with a KMMM using one motif. **B.** The parameters (weights in the PWM) of the recovered motif are plotted against the true parameters. **C.** A 7-mer table constructed from synthetic sequence generated with two 8-base long motifs and logos representing the recovered motifs, Nobox (JASPAR ID MA0125.1, Top) and Sox-17 (JASPAR ID MA0078.1, Bottom). The number of motifs was not preset but was learned during inference. **D.** Comparison of the parameters of the two simultaneously recovered PWMs with the true parameters. Motif logos graphically depict motifs, where the total height of each position represents information at that position.

In both experiments, 10 megabases of synthetic sequence were used and inference was performed by Gibbs sampling. The initial pre-processing step of creating the K-mer table took 1.3 seconds. Inference runtimes were 8 hours for recovering CTCF from an 8-mer and 1.1 hours for recovering the two motif mixture from a 7-mer table on a 2011 MacBook Pro. We obtained similar accuracy using variational inference and stochastic variational inference.

### 6.2 Motifs are found in ChIP-seq peaks and DNAse hypersensitive sites

To assess the ability of KMMMs to succeed in a real motif discovery task, we applied the framework to several freely available sequence sets from the ENCODE consortium including two ChIP-seq experiments and one DNAse-Seq experiment[Encode, 2012].

To assess the performance of KMMMs on identifying motifs from ChIP-Seq, we chose to use the transcription factor NRSF, which has well documented binding motif. We downloaded processed ChIP-seq Broadpeaks in embryonic stem cells (Encode ENCFF001UBT), which consisted of 14,800 peaks with an average size of 350 BP. We constructed an 8-mer table with sequence obtained aligning the peaks to the hg19 assembly of the human genome. We then fit a KMMM with a 3rd order Markov chain background model using 5 motifs by Gibbs sampling. We stopped the sampler after it reached a stable mode of the posterior and present the state of the chain at this point as the recovered motifs (Figure 5). We searched the discovered motifs in the JASPAR motif database [Mathelier *et al*, 2013] and present the closest matches to known human motifs when close matches were found. The motif with the highest information content most closely matches the known NRSF/REST motif (Figure 5).

**Figure 5:**
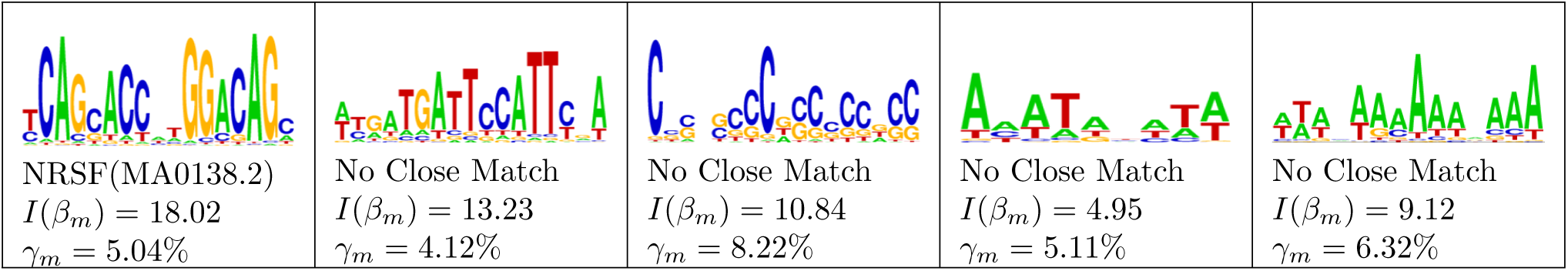
Five motifs found in 30,000 peaks of NRSF ChIP-Seq in human embryonic stem cells (ENCFF001UBT). For each motif, we report the name and ID of the closest match in the JASPAR database along with the motifs Shannon information content, *I*(*β_m_*), and mixing proportion, *γ_m_*. The remaining mixing proportion corresponds to the background component.

We additionally tested the KMMM’s ability to identify motifs from DNAse hypersensitve sites. For this experiment, we looked hypersensitive sites in human lung tissue obtained from the Encode dataset as processed Narrowpeaks (Encode ENCFF474QSW). We constructed an 8-mer table from the 10,000 peaks with the highest signal values, all peaks indexed 150 base-long segments aligned to the GRCh38 assembly of the human genome. We fit a KMMM with 3rd order Markov chain background model with 10 motifs using by Gibbs sampling. We stopped the sampler after it reached a stable posterior mode and present the state of the parameters as the recovered motifs (Figure 6). We then searched for the closest matches present in the JASPAR motif database [Mathelier *et al*, 2013]. Seven of the ten motifs had close matches to known motifs (Figure 6).

**Figure 6:**
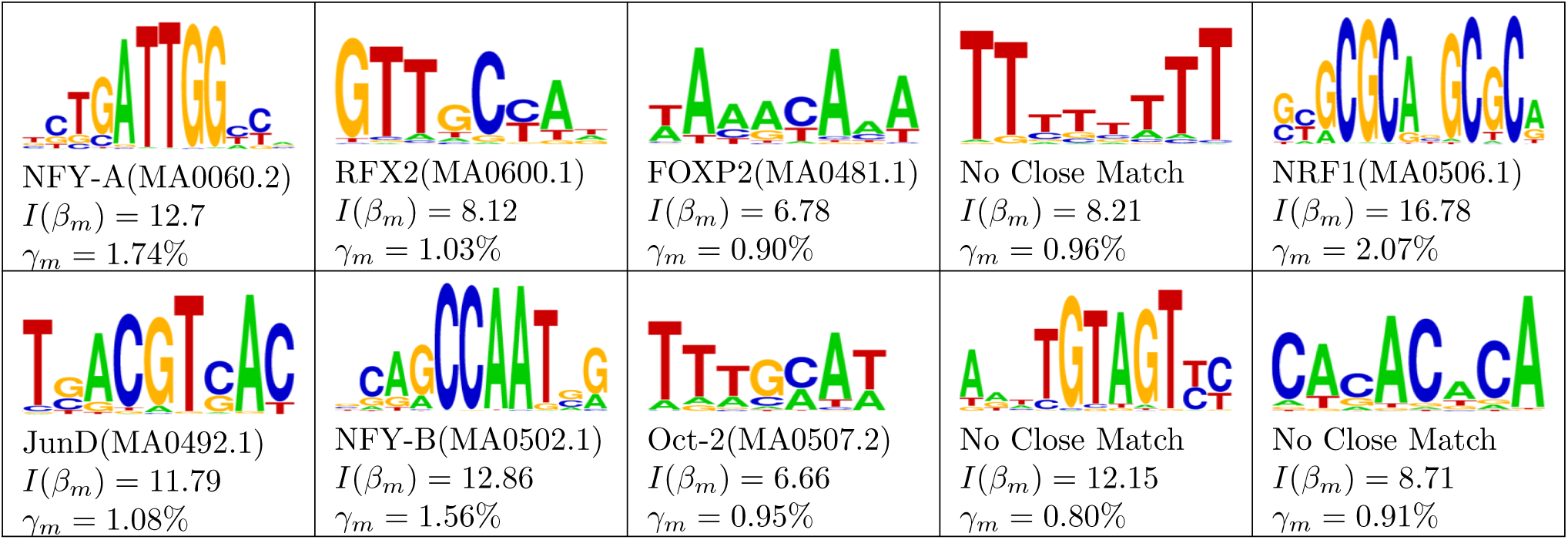
Ten motifs found in DNAse I hypersensitive sites in lung tissue (ENCFF474QSW). Seven of these motifs have close matches with known motifs of Human transcription factors in the JASPAR database. For each motif, we report the name and ID of the closest match in the JASPAR database along with the motifs Shannon information content, *I*(*β_m_*), and mixing proportion, *γ_m_*. The remaining mixing proportion corresponds to the background component.

We additionally applied a KMMM to ChIP-Seq data for the transcription factor MAX. We created a 7-mer table from the processed Narrowpeaks. The sequence set consisted of 46,171 peaks with an average length of 330 BP. As an additional adaptation, we enforced the constraint that the motif be reverse-complement symmetric since MAX is known to bind DNA as a homodimer [Amati *et al*, 1994]. The ability to make this adaptation demonstrates the flexibility of the KMMM framework. We fit this modified KMMM to the 7-mer table, allowing for 8 motifs, by Gibbs sampling for 1000 iterations over approximately three hours. Three of these eight motifs settled into stable modes with high information content (Figure 7). The motif with the greatest information content is the expected E-Box motif (CANNTG) on the left side of Figure 7, known to be bound by MAX [Amati *et al*, 1994].

**Figure 7:**
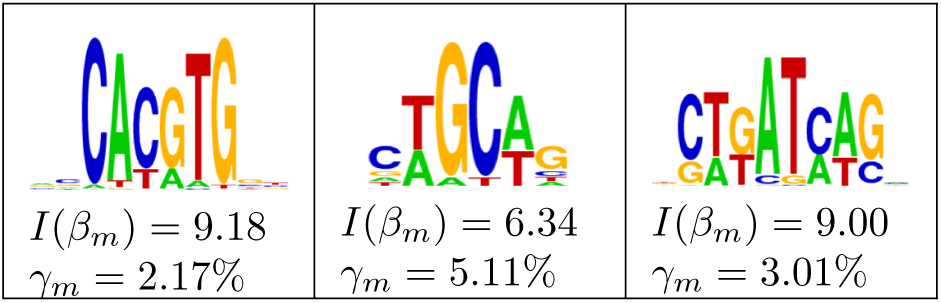
Three symmetric motifs high information content are discovered by fitting a KMMM to 46,171 peaks from ChIP-Seq for the transcription factor MAX. The heights of positions represent the information of the distribution at that position and the relative heights of letters represent the base probabilities. For each motif, we provide Shannon information content, *I*(*β_m_*), and mixing proportion, *γ_m_*. The remaining mixing proportion corresponds to the background component.

### 6.3 Comparison of results with other methods

As KMMMs are generative rather than discriminative models, a fair quantitative comparison of performance to other methods on motif discovery on real genomic sequence is not possible. Nevertheless, a qualitative comparison of motifs found with a KMMM with those found using other method is still informative. To this end, we ran two common alternative methods, HOMER [Heinz *et al*, 2010] and MEME-ChIP [Machanick and Bailey, 2011], on the NRSF ChIP-Seq peaks and DNAse hypersensitive sites that we explored in section 6.2. Comparing the motifs discovered with these methods, we see that some of the motifs are identified by all three of the methods, while others are uniquely identified by each of the algorithms. These results are included and section 12 along with a more detailed discussion. In addition to the comparison of discovered motifs from these real datasets, we also provide results on a synthetic dataset.

## 7 Discussion

We have demonstrated the efficacy of the KMMM for discovering multiple motifs in large sequence sets both on synthetic data and real genomic sequence. However, many of the advantages and limitations of KMMMs have yet to be fully explored.

One advantage of KMMMs arises from the simultaneous inference of multiple motifs. Many approaches to multiple motif discovery identify motifs my iteratively finding new motifs and then erasing from the sequence set to prevent the signal from bleeding over into motifs found subsequently [Bailey and Elkan, 1994]. However, this approach is limited when motifs in sequence set are similar because the signals similar motifs will necessarily interfere. In contrast, KMMMs solve this problem very naturally; when the two similar but not identical motifs are present, the likelihood of a K-mer table will be greater if the motifs are properly modeled. Our results on synthetic data demonstrate this strength. On this task, our inference methods were able to recover the two motifs very closely, despite their similar consensus sequences, which both include “ATTG”. This result stands in contrast to our test of HOMER on this same sequence set, which depending on the background sequence provided, recovered either a single motif, which appeared to be a mixture of the two, or two far less accurate reconstructions (Section 12.3).

Perhaps more significantly, as a Bayesian model class, KMMMs provide many opportunities for extensions within the Bayesian framework. For example, KMMMs can naturally extended to modeling multiple sequence sets that share some of same motifs, but at different levels. This could be achieved by adding additional K-mer tables and associated mixing proportions into the model, but with a single set of motifs. This approach leads to the construction of a topic model and is precisely an instance of latent Dirichlet allocation [Blei *et al*, 2003].

A chief limitation of KMMMs in motif discovery is one inherent to many Bayesian latent variable models; when multiple stable modes within the posterior exist, posterior inference methods will fail to consistently explore the posterior. For example, if Gibbs sampling is used to fit a KMMM with 2 motifs on sequence that contains 3 motifs, it is likely that in a single Markov chain, motif components will fit to only two of these motifs. One method to overcome this problem is to run a Gibbs sampler multiple times to get a better sense of what stable posterior modes may exist.

Another challenge is handling a set of sequences that come from different genomic contexts. Due to the heterogeneity of the genome, different sequences may not be described well by a single background model. This would be the case, for example, for sequences from regions of the genome that have significantly different GC contents. When sequences are pooled together to create a single table, the ability to consider differences in background is lost. One important example of this is poly-A rich areas. In such regions, low motif complexity motifs may have high posterior probability as they are able to better explain background sequence than the Markov chain background model. The poly-T motif discovered in the DNAse-Seq may be an example of this (Figure 6, motif 4).

Despite these limitations, KMMMs is able to discover motifs in real genomic sequence and presents advantages over existing methods when working with motifs in large sequence sets or in more complicated latent variable models. Looking forward, we hope that this model facilitates the development of Bayesian latent variable models that take advantage of the wealth of high-throughput sequencing data available to describe the regulatory state of cells by enabling tractable inference and novel modeling approaches.

## 8 Acknowledgments

Thanks to Dave Blei and the members of the Bussemaker lab, for constructive discussions and guidance.

## 9 Supplement

### 9.1 Notation and terminology

Throughout this report we have used the following terms:

- A sequence is an ordered list of bases, *s* = (*s*_1_, *s*_2_,…, *s_k_*), where each *s_k_* is one of the 4 bases (‘A’, ‘C’, ‘G’, ‘T’).
- A K-mer table, *X*, is a table of counts of length *K* subsequences in the broader sequence, such that *X_s_* is the number of occurrences of sequence, *s*.
- A motif is position weight matrix approximation (PWM). We denote a collection of motifs as *β* = (*β*_1_, *β*_2_,…, *β_M_*), where each *β_m_* = (*β*_*m*,1_, *β*_*m*,2_,…, *β*_*m,K*_) is the set of probabilities over bases for the *K* positions in the motif.
- A background distribution is a normalized distribution over sequences not overlapping with any motifs and is denoted as *ϕ*, where *ϕ_s_* is the probability of drawing *s* from our larger sequence set given that it does not overlap with a motif.
- An offset, denoted as *d*, defines the alignment of a motif relative to a K-mer. More precisely, it is the index of the first position of the motif in the reference frame defined by the K-mer. This is illustrated in Figure 2B. The possible values of *d* are –(*K* – 1), –(*K* – 2),…, (*K* – 1). Given an offset, *d*, we use *s^d^* and 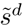 to describe the portions of a sequence attributed to a motif and background, respectively, such that *s* is a concatenation of *s^d^* and 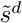. In this notation, 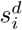 is the *i^th^* base of the overlapping portion of *s*. We use 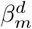 to refer to the portion of motif *m* aligned with *s^d^*, and 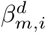 as the distribution over the *i^th^* position in this segment of the motif.
- The background and motif proportions in the underlying sequence are denoted as *γ* = (*γ*_0_, *γ*_1_,…, *γ_M_*). *γ*_0_ is the frequency of background and for *m* > 0, *γ_m_* is the frequency of motif *m*. *γ* lies on the *M*-simplex.

**Algorithm 1.**
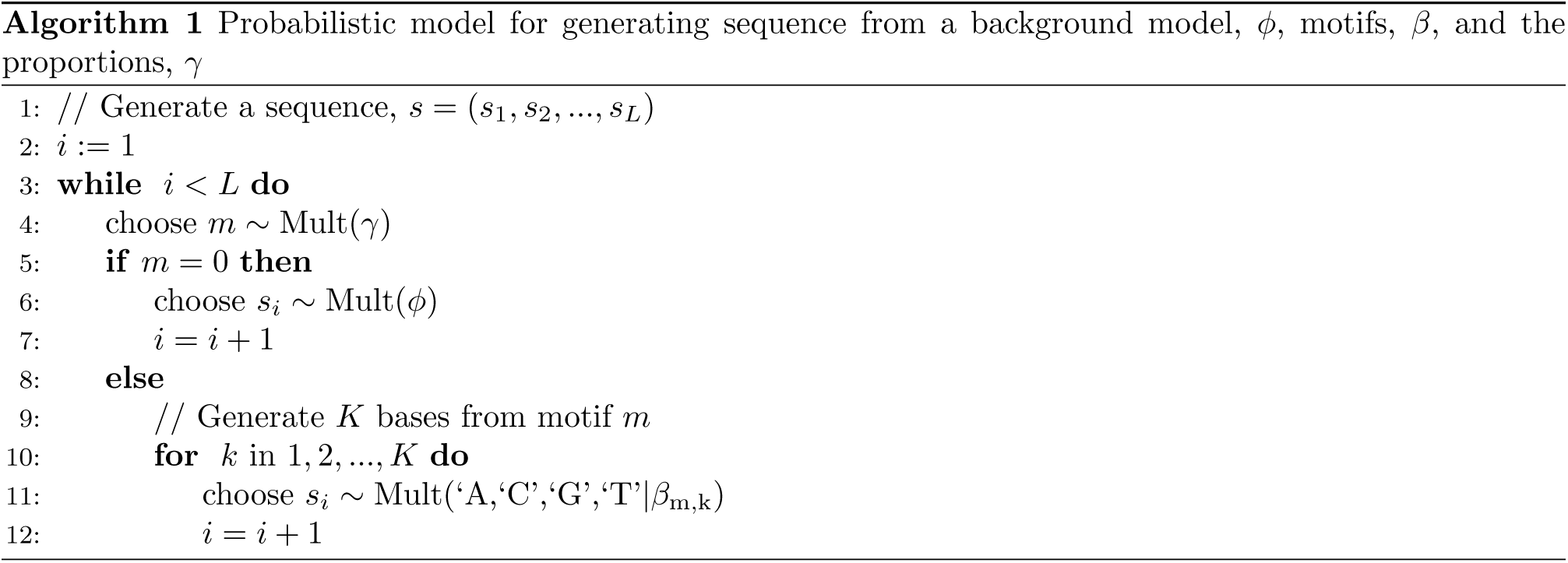
Probabilistic model for generating sequence from a background model, *ϕ*, motifs, *β*, and the proportions,*γ*.

Using this formalism for sequences and motifs, we define a generative model over long sequences bearing latent motifs. This model, presented as Algorithm 1, is an adaptation of the probabilistic segmentation model presented by Bussemaker (2000) in which “stochastic words” are used [Bussemaker et al, 2000, Gupta and Liu, 2011].

## 10 Inference methods

In this section we present details of approximate inference methods for fitting KMMMs. We begin by defining how to compute the counts of bases aligned with motifs from the motif-offset assignment latent variable, *z*, in section 10.1. We then provide details on implementations of Gibbs sampling, variational inference, and stochastic variational inference in sections 10.2, 10.3 and 10.4.

### 10.1 Calculating counts of bases aligned to motifs,η, from motif offset assignments

The counts of each base aligned to each position of motifs must be known in order to calculate the complete conditionals of these latent variables. Since these counts are not directly encoded in the latent variables of KMMMs, we introduced additional notation for each motif, *η_m_*, which refers to the counts of bases from all of the alignments of K-mers against motif *m*. With this notation, *η_m,i,A_* is the number of times we have observed an ‘A’ at position *i* in motif *m*. The value of each component of *η* is defined deterministically by the latent variable *z* through the mapping provided in Algorithm 2.

**Algorithm 2.**
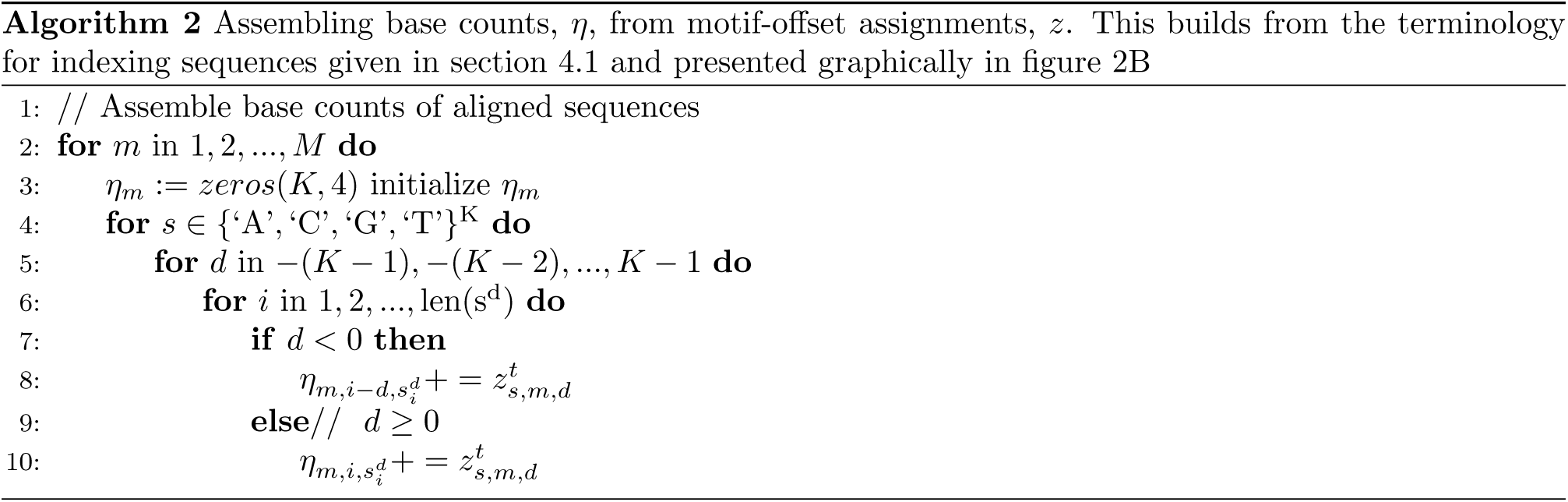
Assembling base counts, *η*, from motif-offset assignments, *z*. This builds from the terminology for indexing sequences given in section 4.1 and presented graphically in figure 2B.

### 10.2 Gibbs sampling

Gibbs sampling is a common method for posterior inference in Bayesian models[?]. To perform inference by Gibbs sampling, we iteratively update of each of a model’s latent variables by sampling from their the complete conditionals. After a period of initialization, often called ‘burn-in’, samples from the posterior are periodically taken by recording the value of the latent variables. Algorithm 3 provides the specific implementation, in which updates are sampled from the complete conditionals established in section 5.1.

**Algorithm 3.**
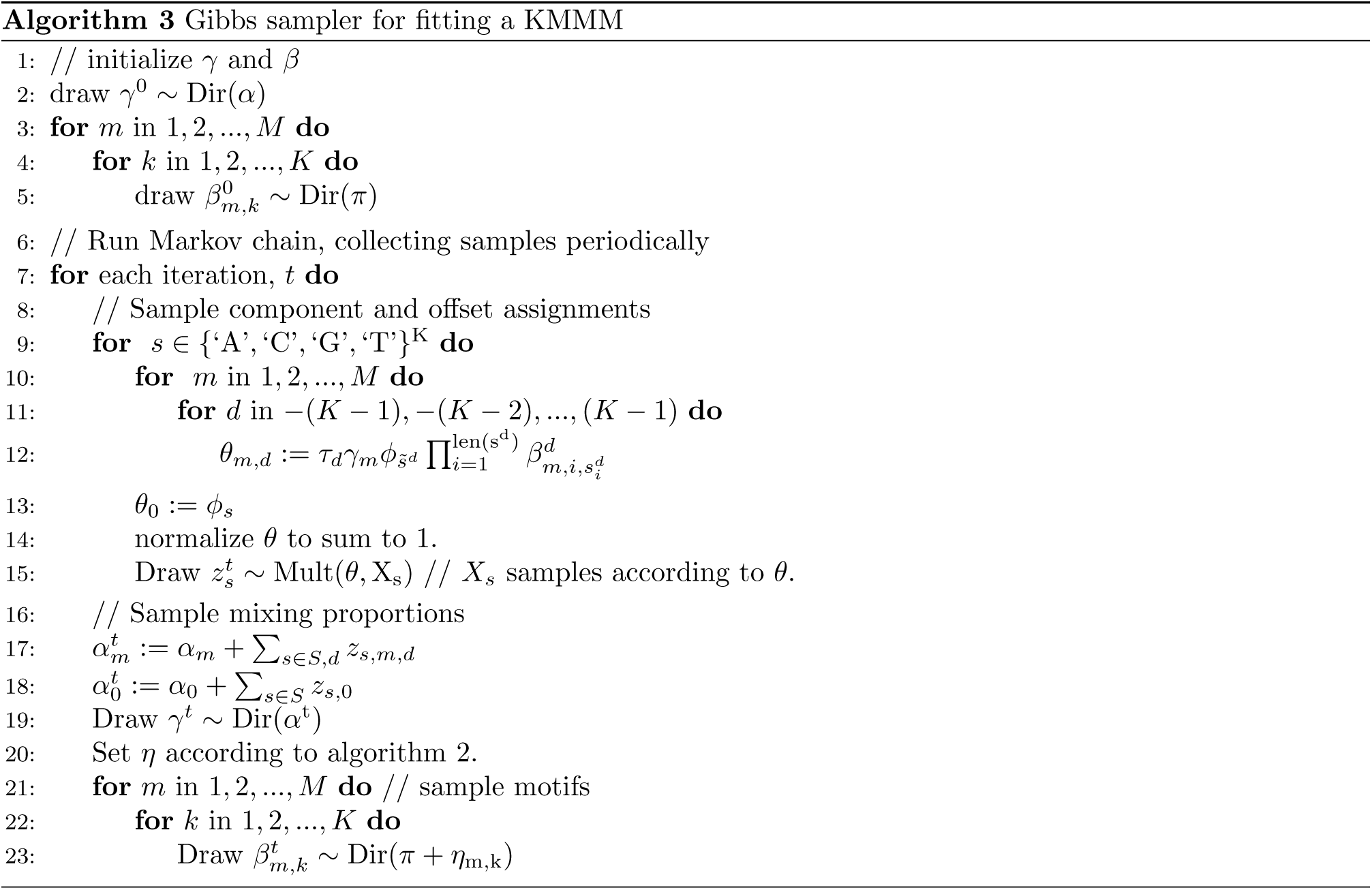
Gibbs sampler for fitting a KMMM.

### 10.3 Variational inference

Variational inference (VI) is a method of inference in which the exact posterior is approximated with a variational distribution, whose parameters are found by optimization[Jordan *et al*, 1999]. The objective of this optimization is minimizing the Kullback-Leibler (KL) divergence between the exact posterior and the variational approximation. In this section we use mean field VI, in which the approximate posterior, *q*, is defined to be a product of conditionally independent distributions over each of the latent variables. As such, the mean field approximation asserts that we can factorize *q* into separate variational distributions:

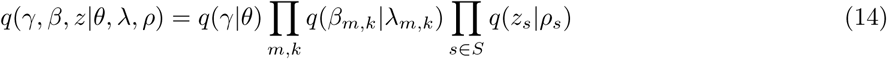

In a conditionally conjugate model, we propose variational distributions in the same family as the complete conditionals of the latent variables they describe[Blei *et al*, 2016]. As such, we use Dirichlet parameters *θ* and λ over *γ* and *β*, respectively, and multinomial parameters, *ρ*, over *z*.

We perform inference by coordinate ascent. We iteratively update our variational parameters to minimize the KL divergence. These updates are take the following analytic forms:

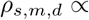

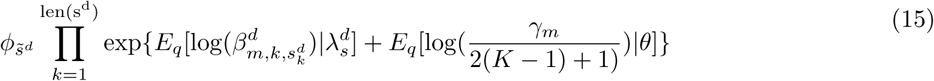

where

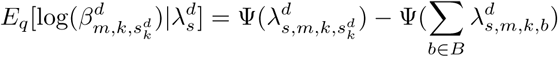

and

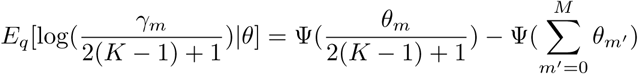

Here, Ψ is the digamma function, which is the derivative of the log of the Gamma function. When performing the update defined by equation 15, we must normalize each *ρ_s_* to 1.

For the variational Dirichlets over the motif PWMs, we use the following update^5^:

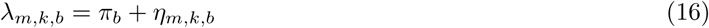

The updates for the dimensions of the variational Dirichlets over the mixing proportions for the motif components are:

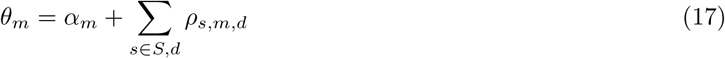

Similarly, for the background component, our update is:

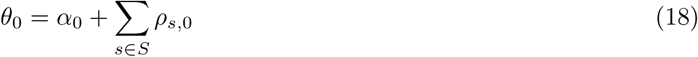

In coordinate ascent VI, we iteratively perform these updates until convergence. The converged variational distributions can then be used to obtain point estimates of the motifs and mixing proportions by taking maximum likelihood values under the approximate posteriors.

In our experiments on the synthetic data with a mixture of two motifs, this VI procedure this converged after roughly 250 iterations in one hour running on a 2011 MacBook and provide faithful reconstructions, *R*^2^ = 0.993.

### 10.4 Stochastic variational inference

Two shortcomings of ordinary VI are the speed of the iterations and the risk of convergence to poor local optima. To increase the efficiency of convergence, and to partially mitigate the risks of arriving at a local optimum, we additionally perform a stochastic implementation of the variational procedure which takes advantage of natural gradients of the exponential family to provide a scalable inference procedure [Hoffman *et al*, 2013]. Here, we use precisely the same variational families as in 10.3, but obtain our estimates at each iteration from only a subset of the K-mers present in the K-mer table. In our experiments we have found success using a forget rate of 0.5, a delay of 10 and a batch size of 1024 K-mers[Hoffman *et al*, 2013].

In our experiments on the synthetic data with a mixture of two motifs, SVI consistently finds the consensus sequences of motifs faster the VI (within approximately 30 minutes) but takes longer than VI to completely converge. We expect that an implementation with more effective parallelization would be more successful.

## 11 Extensions to KMMMs

In this section, we detail extensions to the core KMMM framework which expand its utility in applications motif discovery in real data.

### 11.1 Range of motif models

While we have focused on the PWM model of motifs, several other models exist which can reflect different dependencies and independencies among bases. The KMMM framework can be used with any motif model that can be described by a set of multinomial distributions. This is because we can then assign a Dirichlet prior each of these distributions and maintain the conditional conjugacy necessary for inference in KMMMs.

One alternative motif model incorporates dinucleotide or trinucleotide preferences by describing the sequences that arise from motifs with a 1st or 2nd order Markov chain. For a 1st order Markov chain, we define a motif with start probabilities, *β*^1^, a multinomial distribution over the first base, *s*_1_, and for each of the remaining bases, (*s*_2_, *s*_3_,…, *s_L_*), a matrix of transition probabilities, (*β*^2^,*β*^3^,…,*β^L^*), each defining the probability of each base occurring in that respective position, given the base preceding it. For example, 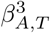 = *p*(*s*_3_ = *T*|*s*_2_ = *A*). This defines the distribution of sequence under the motif as:

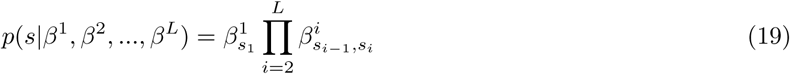

We can describe trinucleotide and even higher order sequence preferences with intuitive extensions of this model using higher order Markov chains, in which transition matrices are replaced with higher dimensional tensors. These can be incorporated into KMMMs because, given a Dirichlet prior over the transition probabilities, Dirichlet-multinomial conjugacy maintains the model’s conditional conjugacy.

A range of other adaptations may also be used including added symmetry constraint, as we showed in our application to motif discovery in MAX ChIP-Seq, and variable length gaps, which can be implemented by introducing gap-size variable as part of K-mer component assignments and a multinomial prior on gap size associated with the gapped motif.

### 11.2 Flexibility of the background model

In our application to synthetic data, we generated sequence with a fully random background model; each base that was not part of a motif was drawn as an ‘A’, ‘C’, ‘T’ or ‘G’ with equal probability, and this simple background model was known and used when inference was performed. However, real genomic sequence is more complicated than this. For example, GC content can vary significantly from 50% and dependencies between neighboring bases can lead to significantly increased frequencies of certain dinucleotides and trinucleotides. One method that can effectively capture low complexity aspects of background sequence is considering the K-mer frequencies in a specific background sequence set [Das and Dai, 2007]. For example, in analysis of ChIP-seq, we often have an negative control (or ‘input’) set, that we want to explicitly compare against. In this context the most effective solution could be to model background sequence according to the frequency of each K-mer in the control set.

In the completely unsupervised case, when no clear negative set is available, using K-mer frequencies is not an option. In this case, one method that effectively can model low complexity aspects of background sequence is a 2nd order Markov chains built from 3-mer counts in the sequence of interest [Bussemaker *et al*, 2000]. While fitting a static background model to an entire sequence data set may be sufficient in some cases, this regime locks some of the signal present in motifs into the background model because some of elevated frequencies of 3-mers that appear in motifs become absorbed into the background model.

To confront this issue, a background model can be learned simultaneously with motifs. We employed this strategy in our application of a KMMM to ChIP-seq peaks and DNAse hypersensitive sites by considering the background model to be an additional latent variable and fit this variable to sequences assigned to the background component during inference. As with alternative motif models, the KMMM framework is compatible with any dynamic background model that can be updated with Dirichlet multinomial conjugacy. This is the case for transition matrices of a Markov chain, which are defined as multinomial distributions upon which one can place Dirichlet priors. With this approach, as we learn to distinguish motifs from background, we reap benefits for being able to distinguish background from motifs; we obtain a more accurate background model and can thereby more accurately infer motifs.

However, using a dynamically updated background model requires keeping track of the sequence attributed to the background component, both the full K-mers and segments attributed to background within motif components, a task that adds a constant factor to the time complexity of inference.

### 11.3 Learning Motif length

Generally when we are interested in discovering a novel motif, we do not know its length *a priori*. However, as we have discussed them so far, KMMMs require a motif to have a well-defined, finite length so that its boundaries may be demarcated in order to explicitly model background sequence. For KMMMs to be practically useful, they must accommodate variable length motifs. In this section we introduce a Bayesian nonparametric extension to KMMMs and inference thereof which enables the length of a motif to be learned as part of inference. This extension makes KMMMs significantly more useful in practice.

To do this, we must integrate the length of the motif into the model as an additional latent variable. We do this by considering a motif to be a countably infinite sequence of distributions, of which only a finite subset of positions govern bases, the bases at all other positions being distributed according to the background model. Prior this section, we described a length L motif as sequence of L distributions, *β* = (*β*_1_,*β*_2_,…,*β_L_*) such that bases aligned with each position of the motif, *η* = (*η*_1_,*η*_2_,…, *η_L_*), were multinomial distributed according to the distribution at that position(Figure 8 left panel).

**Figure 8:**
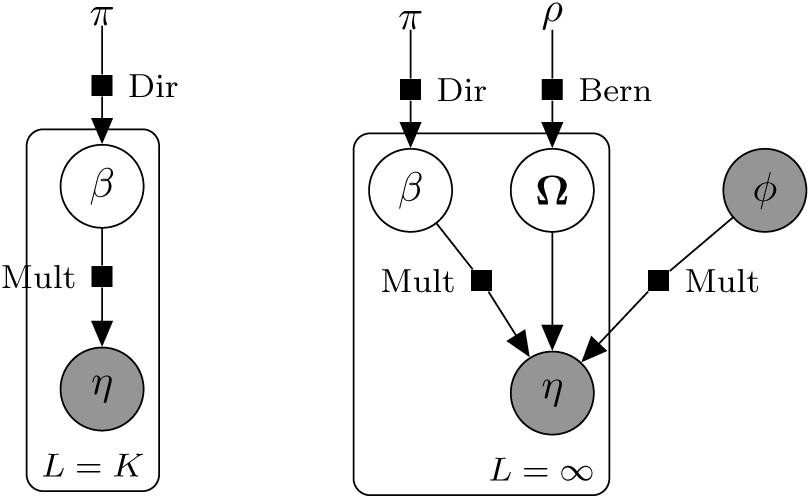
Graphical model representation for PWM motifs in KMMMs. The plate represents multiplicity of the per-position distributions over bases. **Left**: Original motif model has a fixed length, which is equal to the K of the K-mer table. **Right**: Nonparametric extension defines infinite length motifs in which the bases at some positions are drawn from the background distribution.

To accommodate variable length motifs, we extend this model to an infinite sequence of distributions over bases, *β* = (…, *β*_–2_, *β*_–1_, *β*_0_, *β*_1_,…), and add an additional shape random variable, Ω = (…, Ω_–2_, Ω_–1_, Ω_0_, Ω_1_,…), a sequence of binary random variables indicating whether the bases at that position are governed by the motif model or by the background model. Working with the notation for counts of bases aligned to motifs introduced in section 10.1, we can again formulate a generative model and likelihood. If Ω_*k*_ = 1, then we consider the bases at position *k* of the motif, *η_k_* ~ Mult(*β_k_*), else *η_k_* ~ Mult(*ϕ*). Thus, the likelihood of the bases aligned to a given position becomes:

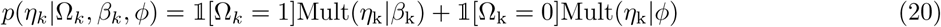

To derive the complete conditional over Ω, we place a prior over it. If we cap the length of the motif at some maximum *L*, we must consider a number of cases that grows exponentially with *L*, which quickly becomes intractable. To circumvent this intractability, we choose a prior over Ω that limits it to a subset of the full space and simplifies inference. First, we constrain motifs to consist of consecutive bases^6^. Second, we propose a truncated geometric prior with parameter, *ρ*, on motif length, where the truncation point is the minimum allowed length for motifs:

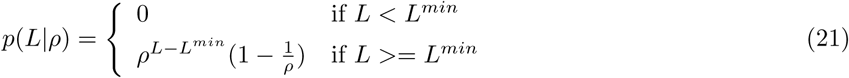

This means that for positions at the edge of the motif (e.g. Ω_0_ and Ω_1_), we can interpret this as a Bernoulli prior, such that Ω_*k*_ ~ Bern(*ρ*) where *k* is the index of the position of either side (start or end) of the motif. The corresponding graphical representation is shown in Figure 8 (right panel).

Now at each iteration, we can sample by motif size by considering, for each side of the motif independently, the posterior probabilities of adding a base, removing a base, or not changing the motif’s length. Under this geometric prior, the relative prior probabilities of these three options are the same regardless of the overall length of the motif. We can now use Bayes’ rule to calculate the complete conditional over the shape variables at the boundary of a motif, Ω_0_ and Ω_1_, where Ω_0_ is the position that could be added and Ω_1_ is the current first position in the motif which could be removed:

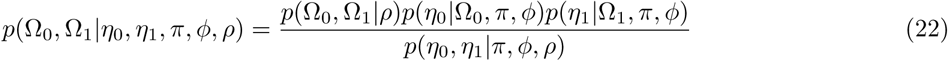

where *p*(Ω_0_, Ω_1_|*ρ*) = *ρ*^Ω_0_+Ω_1_^ (1 – *ρ*)^2–Ω_0_+Ω_1_^. The likelihoods at each position *k*, are:

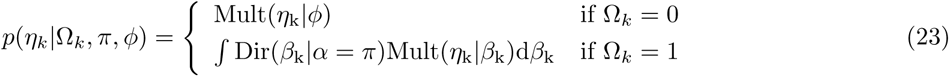

and the normalizing evidence term is calculated by marginalizing over the three cases:

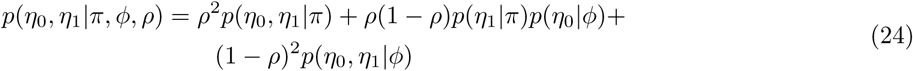

The predictive probability of the multinomial draw given a Dirichlet prior in equation 23 can be calculated with a closed form solution (Tu, 2014):

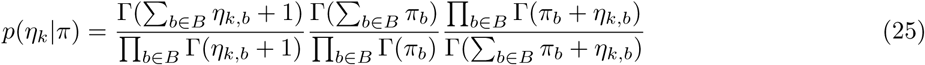

By sampling from this posterior using our Gibbs sampler, we are able to explore the infinite space of motif lengths.

### 11.4 Learning unknown number of motifs

When looking at a set of real genomic sequences, we may not know how many latent motifs exist but still want to identify all that are present. In the context of KMMMs, this question can be rephrased as, how many latent motif components are present in our mixture? One common approach to this type of question is to view it as a model selection problem. As such, one could view the number of motifs as a model specification, and we could pick the best model by trying several different possible numbers of motifs and comparing their performance using a posterior predictive check. However, with this approach we end up re-learning many of the same motifs multiple times across different trials and therefore is not an efficient use of resources. Ideally, we would learn the number of motifs directly from the data as part of fitting the model.

We can accomplish this merging of model selection and fitting by placing a Dirichlet process prior over component assignments, with a base distribution that is a set of uniform Dirichlet priors over the distribution of bases at each position in the motif. This Bayesian nonparametric method provides a prior over all possible partitions of the K-mers across an infinite number of possible motif components.

In order to perform inference in this model, we turn to the Chinese restaurant process (CRP), which defines a generative process for assigning a series of data points across an infinite number of components. According to the CRP, a new data point will be assigned to an existing component with probability proportional to the number of items already assigned to it but could instead be instead be assigned to a yet unseen component with probability proportional to a fixed concentration parameter, *α*_0_. As such, the CRP induces a distribution over partitions of exchangeable items of data [Pitman, 2002]. Under this construction, we formulate a prior over component assignment of any K-mer, *m_i_*, based on the assignments of the every other K-mer, *m*_–*i*_, as:

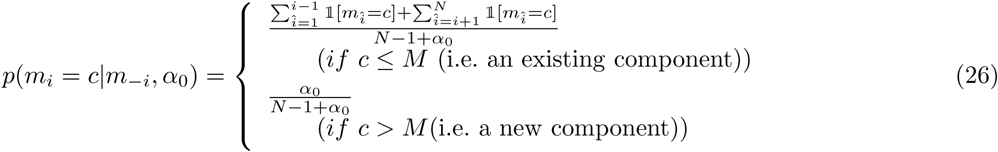

In this way, the nonparametric CRP prior circumvents the need for the explicit mixing proportions latent variable, *γ*. By replacing *γ* with this prior in our formulation of the complete conditional over *z* in equations 9 and 10, we can perform inference over an infinite number of motif components under a Dirichlet process prior using a collapsed Gibbs sampler.

In our application of this technique to synthetic data with two motifs, our Gibbs sampler converged to a stable mode after 1000 iterations. Inference with VI using this extension does not arise as naturally, however.

## 12 Qualitative comparison to other methods

### 12.1 NRSF ChIP-Seq

In order to perform a qualitative comparison between KMMMs and alternative methods for identifying motifs in ChIP-Seq peaks, we ran HOMER and MEME-ChIP on the same NRSF ChIP-Seq experiment, Encode ENCFF001UBT [Encode, 2012]. Both HOMER and MEME-ChIP were able to identify enriched motifs (Figure 9, Figure 12.1).

**Figure 9:**
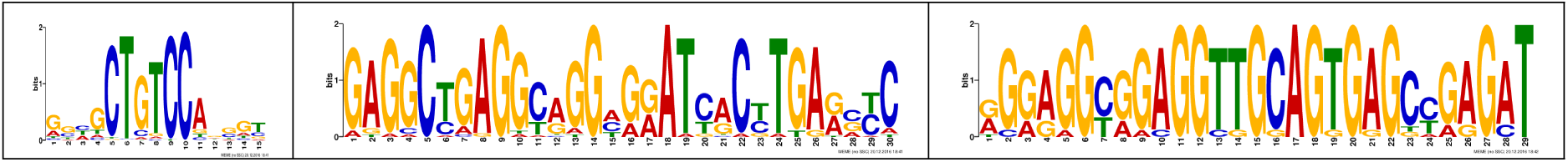
The first three motifs in 30,000 peaks of NRSF ChIP-Seq in human embryonic stem cells (ENCFF001UBT) found using MEME-ChIP.

The first motif identified by MEME-ChIP matches the second half of the motif identified with a KMMM (Figure 9, Figure 5. However, none of the other four motifs identified in our application of a KMMM match either of the other two motifs identified with MEME-ChIP.

**Figure 10:**
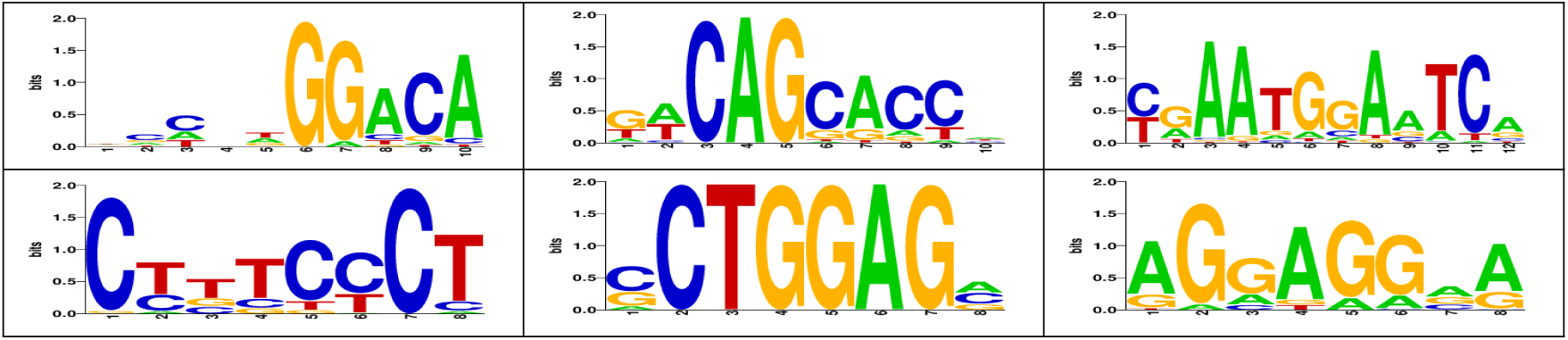
Six motifs in 30,000 peaks of NRSF ChIP-Seq in human embryonic stem cells (ENCFF001UBT) as identified by HOMER.

The first two motifs align partially with segments of the first, long motif motif identified using a KMMM (Figure 12.1, Figure 5). The third motif identified with HOMER matches well to the reverse complement of the second motif identified with a KMMM (Figure 12.1, Figure 5).

### 12.2 DNAse-Seq in human lung tissue

In order to perform a qualitative comparison between KMMMs and alternative methods for identifying motifs in DNAse hypersensitive sites, we ran HOMER and MEME-ChIP on the same DNAse-Seq experiment, Encode ENCFF001UBT [Encode, 2012]. Both HOMER and MEME-ChIP were able to identify enriched motifs (Figure 11, Figure 12). We report the top three motifs found in DNAse hypersensitive sites by MEME-ChIP

**Figure 11:**
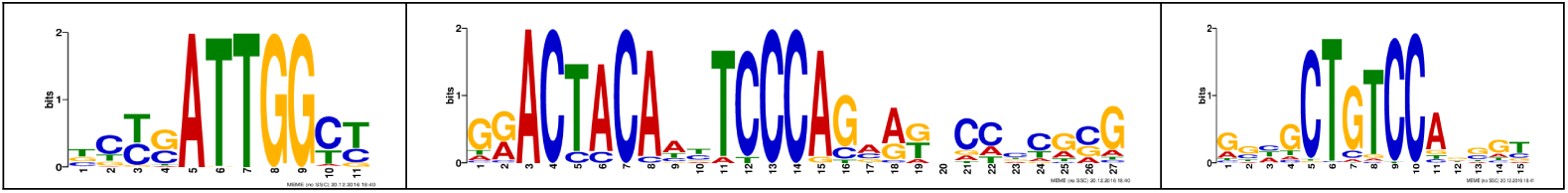
Top three motifs found in DNAse hypersensitive sites in human lung tissue using MEME-ChIP.

The first of the motifs identified by MEME-ChIP, with consensus sequence “ATTGG”, appears to be very similar to motifs 1 and 7 found using a KMMM(Figure 6). This motif also seems to be present in the HOMER results (Figure 12).

**Figure 12:**
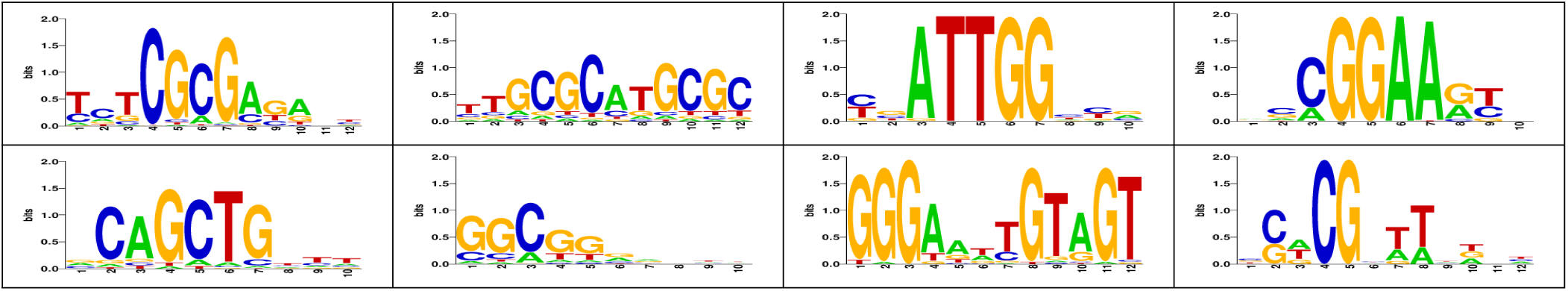
Eight motifs found in DNAse hypersensitive sites using HOMER. We provided the reference genome, hg19, as background sequence set, from which the algorithm picked a subset to use for comparison to peak sequences.

Of the other motifs identified using HOMER, one other also seems to have close matches within the set found in our application of a KMMM to the same sequence set, with consensus “GCGCATGCGC” (Figure 12).

Due to the complexity of DNAse-Seq, which contains a large number of different motifs, one would not expect a naive implementation of any of these methods to return all of the motifs present, so it is not surprising that these three methods do not report the same results. It is encouraging, however that at least two motifs similar to those found by other methods are discovered with a KMMM.

**Figure 13:**
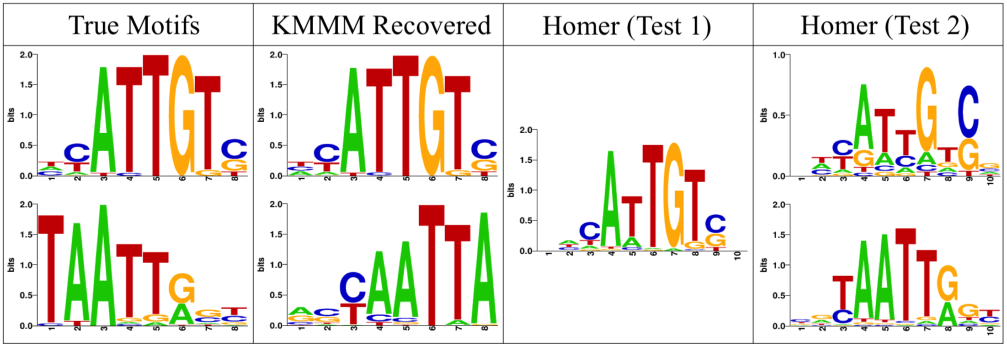
Motifs used to generate sequence and their recoveries. **Col 1** Two 8-base long motifs, Sox-17 (JASPAR ID MA0078.1, Top) and Nobox (JASPAR ID MA0125.1, Bottom), used to generate sequence. **Col 2** The motifs recovered by a KMMM, as presented in Figure 4C. **Col 3** A single motif recovered by running HOMER on the sequence set using a fully random synthetic negative set, reflective to the true background. **Col 4** Two motifs recovered by running HOMER with full human genome provided as a mis-specified background set.

### 12.3 Mixture of motifs in synthetic sequence

As an additional test, we compared KMMMs to HOMER on the synthetic motif discovery task presented in Figure 4C and 4D. To perform this comparison, we ran HOMER on the same sequence set we had used to create the 7-mer table to which we had fit a KMMM. Because HOMER requires a negative set, we tested the tool with two different background sequence sets, obtaining different results under these two conditions. We first ran HOMER with a fully random background set, which was reflective of the true background model used to generate the sequence set. In this test, HOMER discovered only one motif, which appeared to be a composition of the two motifs used, sharing the consensus sequence “ATTG” (Figure 12.3, Col 3). In the second test, we used the entire reference human genome, hg19, as a negative set, from which HOMER picks a subset of sequences [Heinz et al, 2010]. In this test, HOMER successfully identified two motifs, but neither was particularly close to the true motifs used to generate the sequence (Figure 12.3, Col 4).

We believe that this toy example illustrates an advantage of the KMMM framework. By modeling a mixture of multiple motifs simultaneously, the two similar motifs do not interfere with one another as seems to occur when running HOMER naively on this sequence set.

## 13 Practical Notes

### 13.1 IComplexity of inference

The primary bottleneck in Gibbs sampling and coordinate ascent VI is in updating *z* and the variational distribution over *z*. In each iteration, we have *O*(*K* * *K*) computations to score each K-mer with each motif, because we have on the order of *K* positions to compare for roughly *K* possible offsets. The number of K-mers scales as *O*(4^*K*^). We additionally scale linearly in the number of motifs. The total time complexity of these updates is therefore *O*(*K*^2^ * 4^*K*^ * *M*). However, it should be noted that for both variational inference and Gibbs sampling, this update step is easily made parallel. Additionally, one can use efficient stochastic optimization with variational inference by taking advantage of natural gradients of the expectation lower bound with respect to the model’s variational parameters[Hoffman *et al*, 2013].

Notably, the time and space complexity of inference do not depend on the total amount of sequence, as this is reflected only in the magnitude of the entries in the K-mer table. The only step in motif discovery using KMMMs that does scale with the quantity of sequence is the generation of the K-mer table during pre-processing, whose complexity is linear with respect to sequence length.

In the current implementation, we loop through all the possible offsets of all of the motifs for each of the 4^*K*^ K-mers in Python. In the trivial case of the Gibbs sampler identifying 10 motifs of approximately length 10 on an 8-mer table, this means looping through 4^8^ * 10 * (8 + 9) ≈ 1.0 * 10^7^ computations. A re-implementation in a faster language would likely increase the speed of inference significantly.

A second approach is to use intelligent initialization, to reduce the time spent burning in the sampler. Using initialization based on over-represented K-oligomers (given the background model) is a chief method that allows for the high efficiency of existing methods [Heinz *et al*, 2010]. Running a Gibbs sampler or VI with initialization that reflects over-represented K-mers would likely lead to significantly faster convergence.

### 13.2 Hyperparameter selection

As KMMMs are Bayesian models, their priors should be set to reflect beliefs about the values of latent variables. For example, for ChIP-Seq, we can assume approximately one occurrence of motif per peak and use this belief to set our prior on mixing proportions, *α*. In the NRSF data, the average peak size is 350 bases, so if we assume that the motif we are looking for is about 11 bases long and if we are using 8-mers, we expect 18 K-mers to overlap with an occurrence of a motif per peak. This equates to a mixing proportion of about 0.05 for motifs. We use a strong prior on this, so that the data cannot change this. Setting the prior to have roughly as much weight as the data, we can set it to: *α*_0_ = (# of peaks) * (peak – size) and each *α_m_* = 0.05 * (# of peaks) * (peak – size). The Dirichlet prior on motif parameters, π, is less significant; since we do not have a strong prior knowledge about the motifs we would like to find, we use a uniform prior, π = (1,1,1,1).

Another choice that defines a specific implementation of a KMMM is the length of the K-mers. With longer K-mers, motifs are more clearly differentiable from background, which leads motif to be found in a smaller number of iterations. However, this benefit comes at a high computational cost since the time complexity of inference increases exponentially with K. We find 7-mers or 8-mers to be effective in practice.

Parameters must also be chosen to reflect the number of motifs thought to be present in the mixture. In theory, we would hope that the use of the Dirichlet process prior over the mixing proportions, *γ*, introduced in 11.4 would eliminate the need to make this choice. In our experiments with motif in synthetic sequence, we found this method successfully achieved this goal (Figure 4). However, when we applied this method to real data we found it necessary to limit the maximum number of components in the mixture. Without this constraint we found that the number of components would grow to well above 20 motifs. Because the time complexity of each iteration scales linearly with the number of motifs in the mixture, we chose to limit the number of components in order to keep inference fast. We believe that this observation indicates that a many signatures in the sequence exist which are of higher order than could be described by the 3rd order Markov chain used to define our background model.

### 13.3 Reverse complements

Throughout this paper we have nearly completely ignored the issue of reverse complements. In practice we need to consider the fact that if a motif is present, K-mers that align with the motif and its reverse complement will be enriched. Additionally, since we do take orientation information to be meaningful, keeping track of which strand a K-mer is present on while constructing a K-mer table is unnecessary. We take this into account by 1.) counting occurrences of K-mers and their reverse complements together, which reduces the size of the K-mer table by a factor of two and 2.) doubling the number of motif mixture components, so that we have an additional mixture component corresponding to the reverse complement of each motif for each offset.

### 13.4 Dependence of overlapping K-mers

Creating a K-mer table by using a sliding window with which overlapping K-mer are counted causes each base in an occurrence of a motif to be counted ‘K’ times. This over-counting leads to an over-confident estimate of motifs and mixing proportions. In order to account for this, we down-weight the observed counts of bases by a factor of K. This means adapting the specifications of the complete conditionals in equations 11 and 12 to:

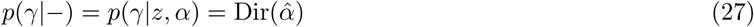

where:

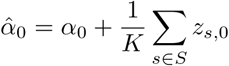

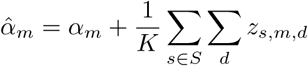

and

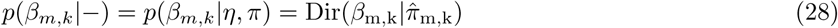

where:

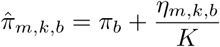

### 13.5 Working with uncertainty

Most current motif methods reason about uncertainty in motifs by first identifying a motif, and then computing statistics related to extent and significance of enrichment. KMMMs approach uncertainty differently; uncertainty in the motifs and their proportions is synonymous with uncertainty in the posterior. As such, uncertainly can be most appropriately modeled by taking into account the uncertainty in the posterior as indicated by the approximate inference methods we have used. Though the correlation structures within the exact posterior are generally very complex in large latent variable models, like KMMMs, in practice, considering the variance within the marginal distributions of the individual latent variables within one of the modes of a posterior can be sufficiently informative. In the context of Gibbs sampling, this means considering the variance in each of the parameters of PWMS across a number of samples from various points within the Markov chain and in VI, variance estimates may be directly obtained from the approximate posterior distributions.

Throughout this paper we have provided only samples from the posterior, in experiments where inference was performed by Gibbs sampling, and point estimates of approximate posteriors from experiments using VI. In our exploration of the variance in motifs found in our reported experiments, we observed that the variance of these parameters was very low, indicating that the posterior has very concentrated density around the point estimates we provided, especially for high information content motifs. We believe, however, that a more rigorous exploration of the uncertainty in motifs discovered with KMMMs, while beyond the scope of this work, could prove to be informative and useful.

## Footnotes

1 Details of this model are provided in section 9.1

2 In a full implementation of a KMMM, reverse complements of sequences must also be marginalized over. This issue is not discussed here, however, because their implementation is a technically intuitive extension of what is discussed in this report.

3 Each offset is equally likely because each occurrence of a motif produces a K-mer with each overlap. This is demonstrated graphically in figure 2B.

4 The notation *p*(*θ*|–) refers to complete conditional over *θ*, i.e. its conditional probability given observed variable, hyperparameters, and all other latent variables

5 This uses the notation for base counts established in Algorithm 2.

6 If Ω_*k*_ = 1 Ω_*k*+1_ = 1 ⇉ Ω = 1.

